# Mitochondrial Genomes Of The Regionally Extinct Nittany Lion (*Puma Concolor* From Pennsylvania)

**DOI:** 10.1101/214510

**Authors:** Maya N. Evanitsky, Richard J. George, Stephen Johnson, Stephanie Dowell, George H. Perry

**Affiliations:** Department of Biochemistry and Molecular Biology, Pennsylvania State University; Department of Anthropology, Pennsylvania State University; U.S. Fish and Wildlife Service, Northeast Fishery Center; Department of Biology, Pennsylvania State University; Huck Institutes of the Life Sciences, Pennsylvania State University

**Author notes:** **Corresponding author:** George H. Perry.

## Abstract

Mountain lions (*Puma concolor*) were once endemic across the United States. The Northeastern population of mountain lions has been largely nonexistent since the early 1800s and was officially declared extinct in 2011. This regionally extinct mountain lion is Pennsylvania State University’s official mascot, where it is referred to as the ‘Nittany Lion’. Our goal in this study was to use recent methodological advances in ancient DNA and massively parallel sequencing to reconstruct complete mitochondrial DNA (mtDNA) genomes of multiple Nittany Lions by sampling from preserved skins. This effort is part of a broader Nittany Lion Genome project intended to involve undergraduates in ancient DNA and bioinformatics research and to engage the broader Penn State community in discussions about conservation biology and extinction. Complete mtDNA genome sequences were obtained from five individuals. When compared to previously published sequences, Nittany Lions are not more similar to each other than to individuals from the Western U.S. and Florida. Supporting previous findings, North American mountain lions overall were more closely related to each other than to those from South America and had lower genetic diversity. This result emphasizes the importance of continued conservation in the Western U.S. and Florida to prevent further regional extinctions.

## Introduction

The *Puma concolor* (mountain lions; also referred to as cougars, pumas, panthers, or catamounts) lineage diverged 3.19 MYA from that of its closest relative, *Miracinonyx trumani*^1^. Mountain lions were widespread across both North and South America by at least 2 MYA until ~13,800-11,400 BP (years before present) when a Late Pleistocene extinction event eliminated up to ~80% of all large mammals from North America, including mountain lions^2^. North America was subsequently re-colonized by immigrant populations of South American *P. concolor* 12,000-10,000 years ago^3,4^.

From that time until recently, *Puma concolor* was endemic throughout much of the Americas^5–7^. However, following European colonization, North American mountain lions were subject to targeted hunting by farmers who perceived them as threats to their livestock and settlements. Simultaneously, land clearing for farming and grazing led to the loss and fragmentation of many forests and to the significant depletion of many deer populations, a primary prey resource for mountain lions^6–8^.

As a result, today in North America mountain lion populations can only be found in Mexico, in the Western United States and Canada, and in the southern tip of Florida^3,9^. In the Northeastern United States, mountain lions have been almost nonexistent since the early 1800s^6^. The existing populations in the Western U.S. and Florida are at risk of extinction due to continued habitat fragmentation and encroachment from suburban and urban sprawl^10,11^. The last documented observation of *P. concolor* in Pennsylvania was in 1874, and the last *in situ* mountain lion known to exist in the Northeastern United States was killed in 1938 in Maine^12^.

The eastern mountain lion was officially declared regionally extinct in 2011 by the U.S. Fish and Wildlife Service^7^. Very recently, there have been many mountain lion sightings in the Northeastern United States (multiple confirmed), all of which reflect long-range individual migrants and the recent range expansion of Western mountain lions, rather than *in situ* recovery of a surviving endemic Northeastern United States mountain lion population^13,14^.

As many as thirty mountain lion subspecies have been described, including thirteen located north of Mexico^15^. Three of these ‘classical’ North American subspecies would be considered Endangered (including the Florida panther *P. conocolor coryi)*, while the Eastern cougar (*P. concolor couguar)* and the Wisconsin puma (*P. concolor schorgeri*) are extinct^3^. However, in a genetic study, Culver et al.^3^ sequenced up to 891 bp of the mitochondrial genome and genotyped up to 10 nuclear genome short tandem repeat (microsatellite) loci from 186 North American mountain lion individuals, including three museum specimens from the Northeastern United States. Their analysis failed to support the classification of multiple North American subspecies, with nearly all individuals from North America, including the three Northeastern United States mountain lions having an identical mitochondrial DNA (mtDNA) haplotype. Questions remain concerning fine-scale patterns of mountain lion genetic diversity when considering the complete mtDNA genome (17,153 bp)^16^ and/or large numbers of single nucleotide polymorphism markers from the nuclear genome.

Our goal is to raise awareness of species conservation and extinction risk issues among the students of Pennsylvania State University and the broader Penn State community, for whom the ‘Nittany Lion’ is a beloved mascot, via the application of recent ancient DNA (aDNA) and genomic sequencing advances to preserved skin samples from nineteenth century Pennsylvania mountain lions and then by disseminating our results. This first study focused on reconstructing complete mtDNA sequences from Northeastern mountain lion specimens curated in Pennsylvania museums and private collections that had been preserved in the late 1800s via taxidermy (**Figure 1**). We focused initially on mtDNA because there are typically hundreds to thousands more mtDNA genome copies than nuclear genome copies per eukaryotic cell, including for skin cells^17^. Thus, for aDNA applications, when DNA quantity and quality is limited (and especially in this case, in which we are working with samples upwards of 150 years old exposed to taxidermy processes that substantially diminish DNA quality^18^), mtDNA is a typical first step. This initial analysis provides information about the quality of endogenous DNA in general in these samples, setting the stage for subsequent complete nuclear genome sequencing of the best preserved samples to facilitate more powerful analyses of mountain lion genetic diversity and phylogeography.

**Figure 1:**
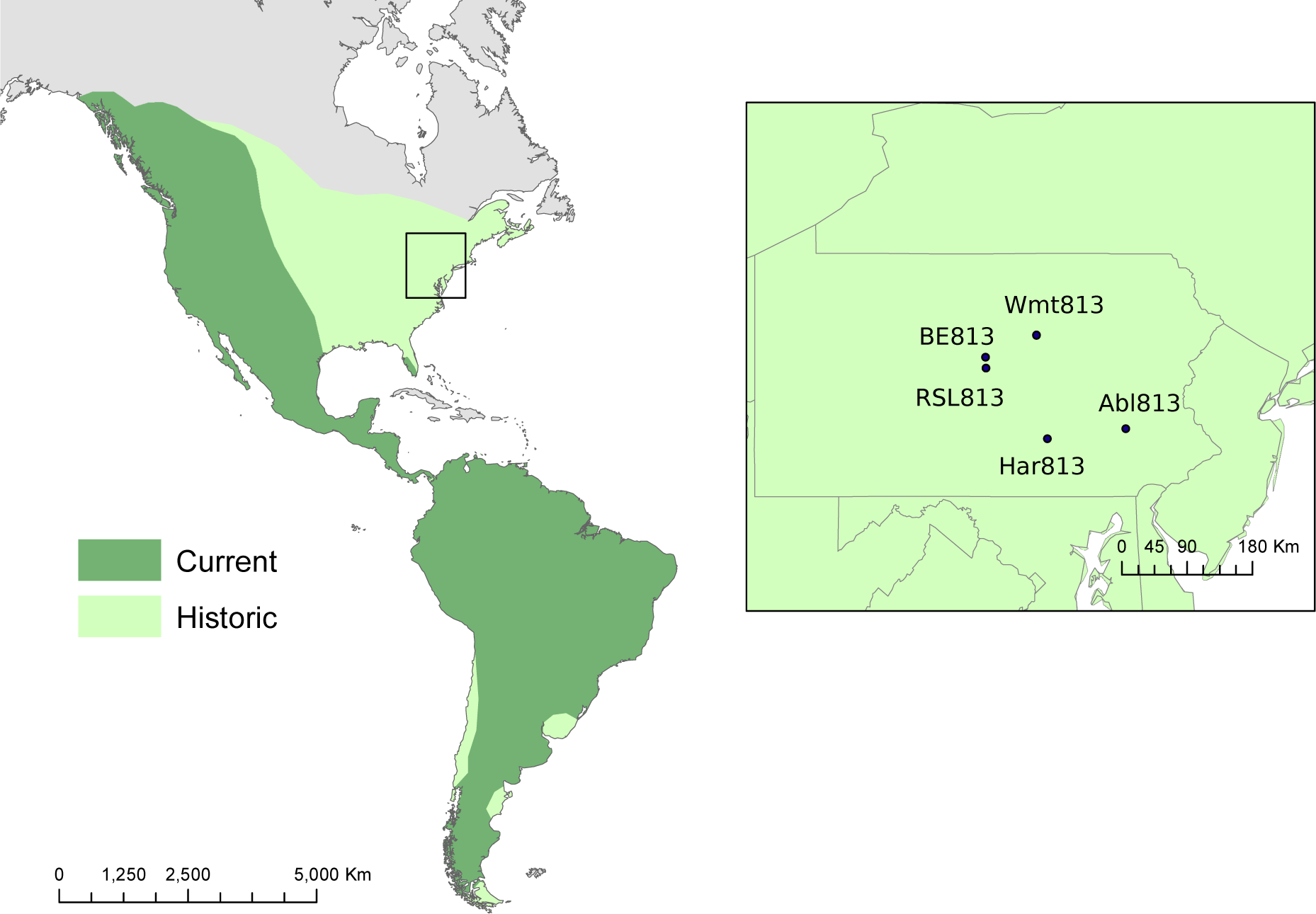
Map of Collection Sites and Mountain Lion Ranges. Samples were collected from Albright College; Bald Eagle State Park; the Pennsylvania State Museum; Howard, PA; and the Thomas T. Taber Museum. The current and historic ranges for mountain lions are also displayed^3^. Map source: ESRI. The map was generated using ArcGIS Pro 1.2 by ESRI (available at: http://www.esri.com/).

## Methods

### Sample Collection

Skin samples were obtained from six Northeastern mountain lions preserved using taxidermy prior to the 1900s (**Figure 2**). Due to the age and morphological importance of the specimens, samples were collected with as little damage to the specimen as possible. A lab coat, face mask, and gloves were worn to limit modern DNA contamination, as well as to provide personal protection from any harmful chemicals used in the preservation process. Five samples were personally collected by M.N.E. using sterilized scalpel and scissors from local Pennsylvania museums, the visitor center at Bald Eagle State Park, and a private residence. The sixth sample was donated by Albright College (**Table 1**).

**Figure 2:**
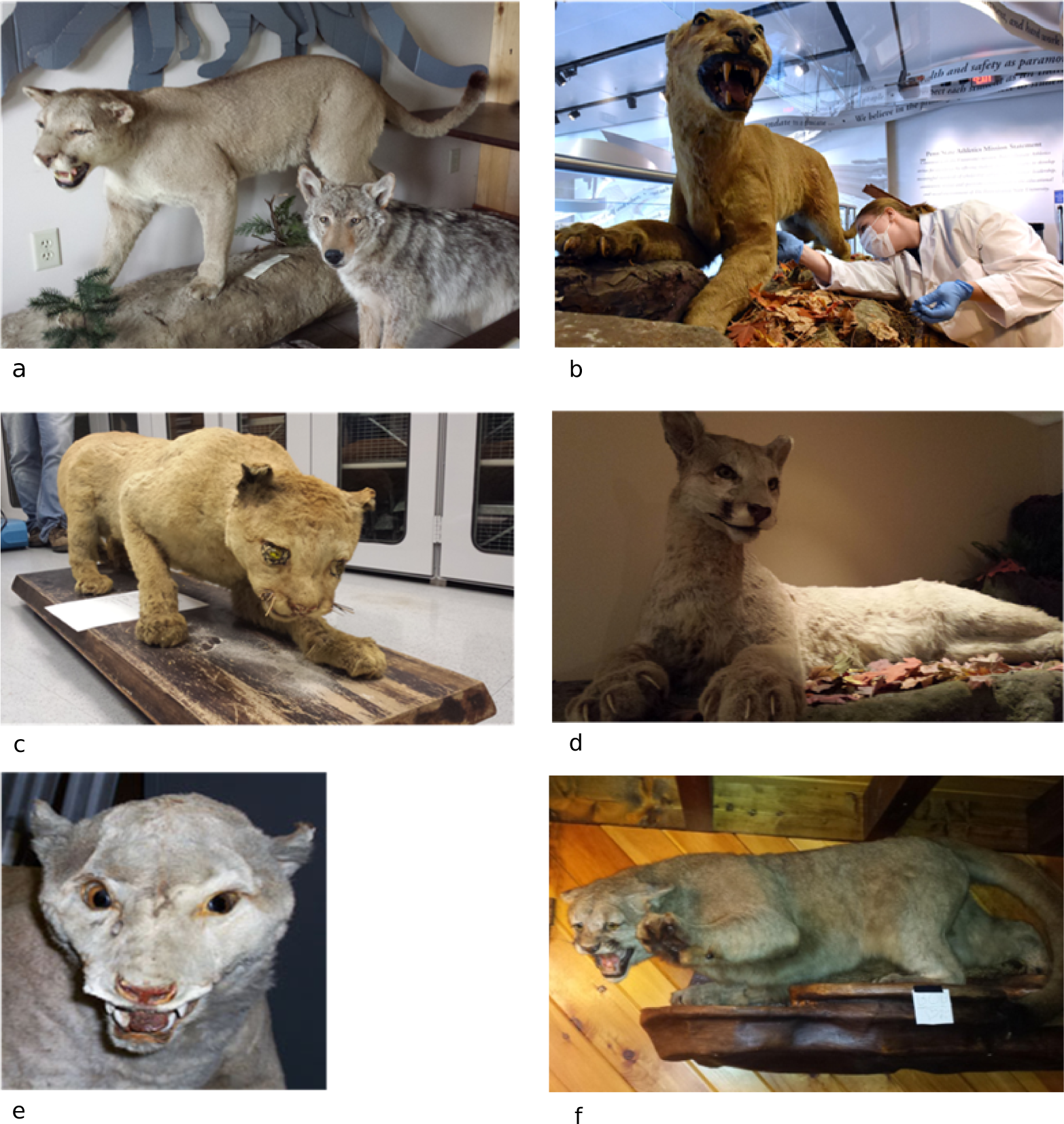
Mountain Lion Specimens and Sampling. The six museum specimens that were sampled for DNA extraction are shown. a) Bald Eagle State Park b) Penn State All-Sports Museum; Photo: Pat Little c) Thomas T. Taber Museum d) Pennsylvania State Museum e) Albright College; Photo: Stephen Mech, Albright College Biology Department f) Rock Spar Lodge. All photos were taken by the authors unless indicated otherwise.

**Table 1:** **Sample Information**

| Sample ID | Current Location | Museum/Owner | Time of Death |
| --- | --- | --- | --- |
| Alb813 | Reading, PA | Albright College | 1870s |
| BE813 | Howard, PA | Bald Eagle State Park Visitor's Center | unknown |
| Har813 | Harrisburg, PA | Pennsylvania State Museum | 1874 |
| RSL813 | Howard, PA | Privately Owned | unknown |
| Wmt813 | Williamsport, PA | Thomas T. Taber Museum | 1880s |
| NL813 | State College, PA | Penn State All-Sports Museum | 1856 |

### DNA Extraction

All sample processing and DNA extractions were performed in a clean lab dedicated to aDNA with standard aDNA protocols^19^. The Penn State ancient DNA laboratory is located in the physics building on campus; the lab is completely separated from any ongoing modern genetics work, and has positive pressure, HEPA filtration, and stringent work protocols to prevent any modern DNA contamination. Samples were digested and the DNA was extracted using a phenol:chloroform protocol modified from de Moraes-Barros & Morgante^20^(described below). All equipment and reagents (as appropriate) were sterilized by UV-irradation for one hour prior to use.

In preparation for digestion, samples were cut into small pieces of approximately 2-3 cm^2^ each. Samples were first immersed in 50% bleach for 30 seconds to remove any surface contaminant DNA, then rinsed with ethanol and molecular grade water three times each. Each sample was washed three times with 1.5 mL of NTE solution (50 mMtris, pH 9; 20 mM EDTA; 10 mMNaCl) to remove potential inhibitors. Samples were then hydrated by incubation for 24 hours in 1 mL of TE solution (tris 10 mM, EDTA 1 mM, pH 7.6) and washed with 70% (w/v) ethanol and molecular grade water before a second hydration in TE solution for a further 24 hours. Samples were cut in half prior to digestion using a sterile razor blade.

For the digestion, one extraction blank was used for every six samples. 500 µL lysis buffer (TrisHCl 10 mM, pH 8; NaCl 400 mM; EDTA 2 mM, pH 8.0; SDS 1%), 100 µL of Proteinase K (20mg/mL) and 100 µL of 500 mM Dithiothreitol were added to each sample. Additional lysis buffer was added until the total volume reached 1 mL. Sample tubes were then placed on a rotary mixer and rotated overnight at 40 rpm with incubation at 55°C or until sample was fully digested. This sample processing and digestion protocol was adapted from Casas-Marce et al.^18^ and de Moraes-Barros & Morgante^20^.

One volume phenol was added to each sample, which was then rotated at room temperature for 5 minutes at 40 rpm and centrifuged at maximum speed (12,900 x g) for 8 minutes until phases separated. The aqueous layer was transferred to a new tube. This process was done twice. One volume chloroform was added to the remaining aqueous layer and rotated at room temperature for five minutes. The aqueous layer was transferred to a clean 1.5 mL tube.

Following phenol: chloroform extraction, the DNA was purified and concentrated using a MinElute PCR Purification Kit with modifications to retain short DNA fragments, modified from the procedure by Dabney et al^21^. Specifically, 6 mL of binding buffer (5 M guanidine hydrochloride, 90 mM sodium acetate, 40% isopropanol, 0.05% Tween-20) was mixed with each sample. Qiagen tube extenders were attached to each MinElute column and placed in 50 mL conical tube. Tube extenders and conical tubes were UV-irradiated prior to use. The extraction-binding buffer mixture was added to a tube extender and centrifuged for four minutes at 1,500 x g in a large centrifuge. The filtrate and tube extenders were discarded and the MinElute column placed in a 2 mL collection tube. The column was dry spun at 3,300 x g in a bench centrifuge for one minute. The column was washed twice with 700 µL PE buffer and spun at 3,300 x g in a bench centrifuge for one minute. The column was spun once more for one minute at maximum speed to remove all traces of ethanol, and then was transferred to a clean 1.5 mL tube. 30 µL of TET was added and incubated on the column for 15 minutes at 37°C. The column was centrifuged for one minute at maximum speed to elute.

### Library Preparation

Genomic sequencing libraries were constructed according to the Meyer & Kircher^22^ protocol with two modifications for aDNA. In modification one, samples were not sheared prior to end repair, as the aDNA samples are already heavily degraded and fragmented^19^. For modification two, in the final PCR amplification step, the samples were divided into two equal parts for duplicate 50 µl reactions. To briefly describe the library preparation process, first any double-stranded DNA fragments with overhanging ends as a result of the aDNA fragmentation process are repaired. Then, oligonucleotide adaptors are ligated to both ends of each DNA fragment that facilitate the massively parallel sequencing process along with the addition of sample-specific “barcode” sequence indices to each fragment. These barcodes make it possible to sequence multiple samples on the same sequencing run (‘multiplex’ sequencing), as sequence reads from each sample can still be analyzed separately based on their unique barcodes.

### MiSeq Test Run

To test the success of extraction and library construction, all samples with concentrations above 15 ng/µl were pooled to equivalent DNA masses (40ng each) and sequenced on the Illumina MiSeq using paired-end, 2x75 bp read lengths. Sequence data were first processed and adaptors removed using a perl script adapted from Martin Kircher designed for aDNA analyses^23,24^. Using this script, we removed adapter sequences and merged forward and reverse reads with minimum 11 nt overlap and combined phred quality score of merged sites greater than 20. Sequence reads were then aligned to the complete *Puma concolor* mitochondrial genome available on Genbank (accession number NC_016470)^16^ using the Burrows-Wheeler Aligner (BWA)^25^. Samples were processed and potential PCR duplicate reads removed using the SAMtools package version 1.2 (using htslib 1.2.1) with default settings^26^. Experimental quality was assessed by quantifying the numbers of total mapped reads and ‘uniquely mapped’ reads following removal of PCR duplicates, the average sequence coverage across the mtDNA genome, and the read length distribution. A uniquely mapped read is one that mapped to the reference genome in only one position.

The C-to-T mismatch distribution was also determined using mapDamage 2.0^27^ and visualized in R Studio^28^. As part of the aDNA degradation process, cytosine residues at the single-stranded fragment ends may deaminate to uracils, which then may be read as a thymine in sequencing, resulting in a higher proportion of C-to-T mismatches and G-to-A mismatches at the 5’ and 3’ ends of DNA fragments, respectively^19^. If this pattern is present, then downstream analyses must account for nucleotides representing potential sequence damage (e.g., by hard-masking to “N” all potentially damaged nucleotides at the first and last ~10 bp of each sequence read, depending on the observed pattern of damage) to limit the chance that this damage would result in incorrect sequence reconstruction. For aDNA samples, this damage pattern tends to increase with sample age^29^. We observed very low proportions of such mismatches for our samples, likely due to their relatively recent ages compared to many aDNA samples (**Supplemental Figure 1**). Damage masking correction was thus unnecessary.

Based on the MiSeq data alone, one of our six samples was successfully sequenced to high coverage (with each mtDNA nucleotide position covered by an average of more than 30 sequence reads), two were sequenced to low coverage (3-5 fold average coverage), and two other samples had evidence of mountain lion DNA present but at levels insufficient for reconstructing the complete mtDNA genome (**Table 2**). For one sample, NL813, we obtained zero sequence reads that could be assigned confidently to the mountain lion mtDNA genome. For this sample, we performed an additional DNA extraction, with the addition of a Microcon 30 kDa centrifuge tube filtration step, and then constructed a new library using the methods described above, before proceeding to DNA capture and Illumina HiSeq analysis (described below), in an attempt to remove any potential arsenic residues remaining from the taxidermy process that may have impeded our initial library preparation.

**Table 2:** **MiSeq Sequence Data Quality Analysis**

| Sample ID | Total Reads | Non-redundant Mapped Reads<sup>*a</sup> | Average Depth Coverage* |
| --- | --- | --- | --- |
| **Alb813** | 3698194 | 25 | 0.07 |
| **BE813** | 9381398 | 6705 | 32.1 |
| **Har813** | 5164026 | 71 | 0.22 |
| **NL813** | 402 | 0 | 0 |
| **RSL813** | 680853 | 1178 | 4.58 |
| **Wmt813** | 1722516 | 977 | 3.15 |

\*Calculated after removing potential PCR duplicate reads. All reads were mapped to the *P. concolor* reference mitochondrial genome<sup>16</sup>.
<sup>a</sup>Reads that mapped to the reference mtDNA genome at only one position.

### DNA Capture

For five of our six samples, we used a DNA capture process to enrich our shotgun libraries for mountain lion mtDNA fragments prior to sequencing on the Illumina HiSeq. Enrichment was unnecessary for sample BE813 due to the high sequencing quality obtained on the MiSeq test run (**Table 2**). For enrichment we used a custom MYcroarray Mybaits kit, which uses biotinylated RNA oligonucleotides designed to bind targeted endogenous DNA fragments within the shotgun library, with the RNA probes designed to be complementary to the DNA sequence of interest^30^. The probes are hybridized to the library and then the product is mixed with streptavidin-coated magnetic beads. Streptavidin forms a strong bond with biotin. Thus, the hybridized, endogenous mtDNA fragments can then be separated from the remainder of the shotgun DNA library with a magnet and repeated washes, resulting in enrichment for the targeted regions of interest.

Our biotinylated RNA probes were designed using the *P. concolor* reference mitochondrial genome^16^ (Mybaits probeset #150601). Mybaits protocol version 3.0 was followed for enrichment, using the manufacturer’s suggested modifications for aDNA. Specifically, baits and DNA sequencing libraries were incubated together for 40 hours, rather than the standard 16-24 hours suggested for modern DNA. Following binding of the hybridized DNA-RNA fragments to the beads, 180µl of wash buffer was used to dilute non-targeted fragments in a series of four total washes. The captured libraries were amplified with 24 PCR cycles.

### Illumina HiSeq sequencing

All six samples (the five enriched and one non-enriched libraries) were sequenced in parallel on a single lane of the Illumina HiSeq 2500 at the Penn State Huck Institutes of the Life Sciences Genomics Core Laboratory, with paired-end 2x75 bp reads. Samples were analyzed for quality according to the same protocol as described above. Five out of the six samples were successfully sequenced. One sample (NL813) failed on both the MiSeq and HiSeq runs, indicating either the extraction failed or the arsenic reported to be in the sample interfered significantly with library construction (**Table 3**). All MiSeq and HiSeq read data are available via the NCBI Sequence Read Archive (SRA) under accession number SRP090336. Consensus mtDNA genome sequences for each sample were constructed using the SAMtools mpileup command with the parameter -C50 to adjust the mapping quality of reads with multiple mismatches and the parameter -q20 to impose a minimum mapping quality of 20. Sequence data from both the MiSeq and HiSeq runs were merged using the SAMtools merge command with default settings prior to consensus calling. We required minimums of 2X non-redundant sequence coverage and 80% site identity per site for consensus base calling (**Supplemental Table 1**). Our final consensus sequences for each sample and intermediate processing files from the read alignment are available through Penn State’s Scholar Sphere resource at: https://doi.org/10.18113/S19G6D

**Table 3:**
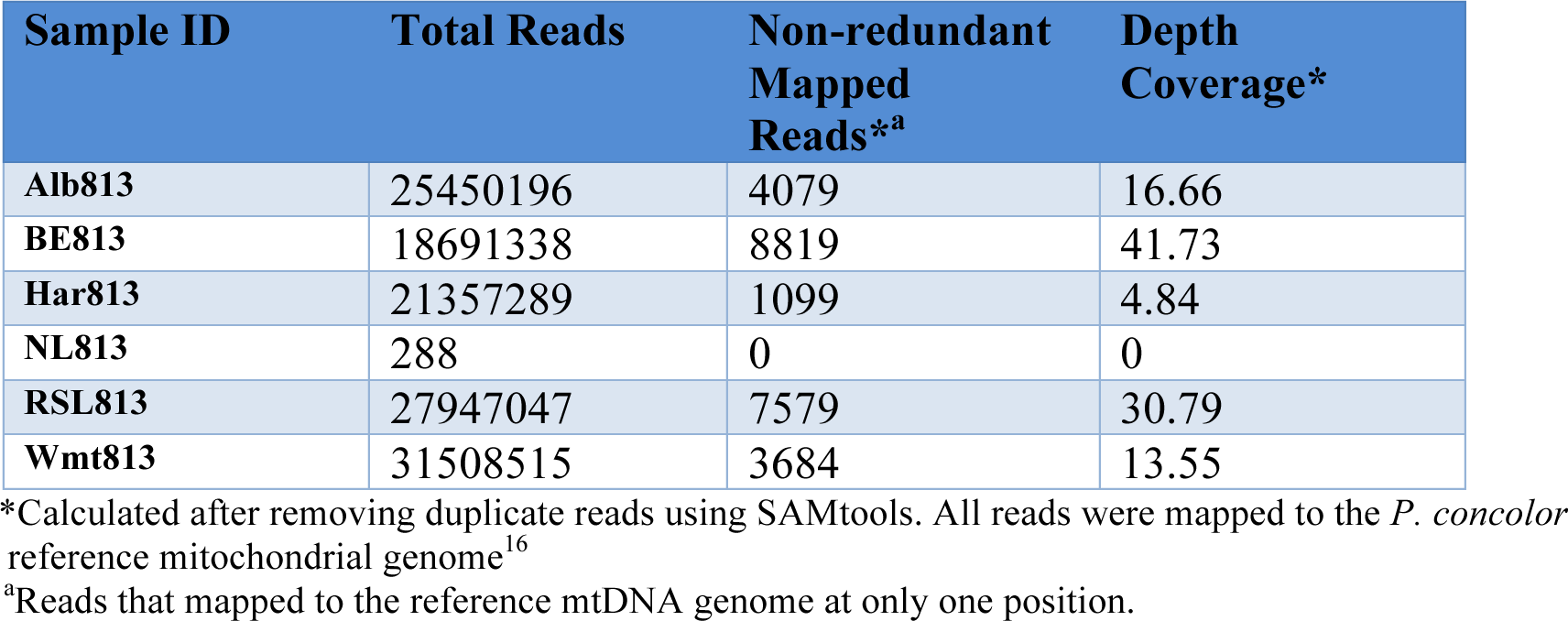
**HiSeq Sequencing Data Analysis**

| Sample ID | Total Reads | Non-redundant Mapped Reads<sup>*a</sup> | Depth Coverage* |
| --- | --- | --- | --- |
| **Alb813** | 25450196 | 4079 | 16.66 |
| **BE813** | 18691338 | 8819 | 41.73 |
| **Har813** | 21357289 | 1099 | 4.84 |
| **NL813** | 288 | 0 | 0 |
| **RSL813** | 27947047 | 7579 | 30.79 |
| **Wmt813** | 31508515 | 3684 | 13.55 |

\*Calculated after removing duplicate reads using SAMtools. All reads were mapped to the *P. concolor* reference mitochondrial genome<sup>16</sup>.
<sup>a</sup>Reads that mapped to the reference mtDNA genome at only one position.

All positions at which the Northeastern mountain lions differed from the *P. concolor* reference sequence were verified visually by examining the depth of coverage and fraction of reads that supported the base substitution at each position where a SNP occurred (see **Figure 3** for one example region containing three SNPs). Each SNP variant call was indeed supported by nearly all reads; instances in which a small minority of reads are different from the consensus likely represent sequencing errors or environmental DNA damage and did not affect the consensus sequences used in our analyses.

**Figure 3:**
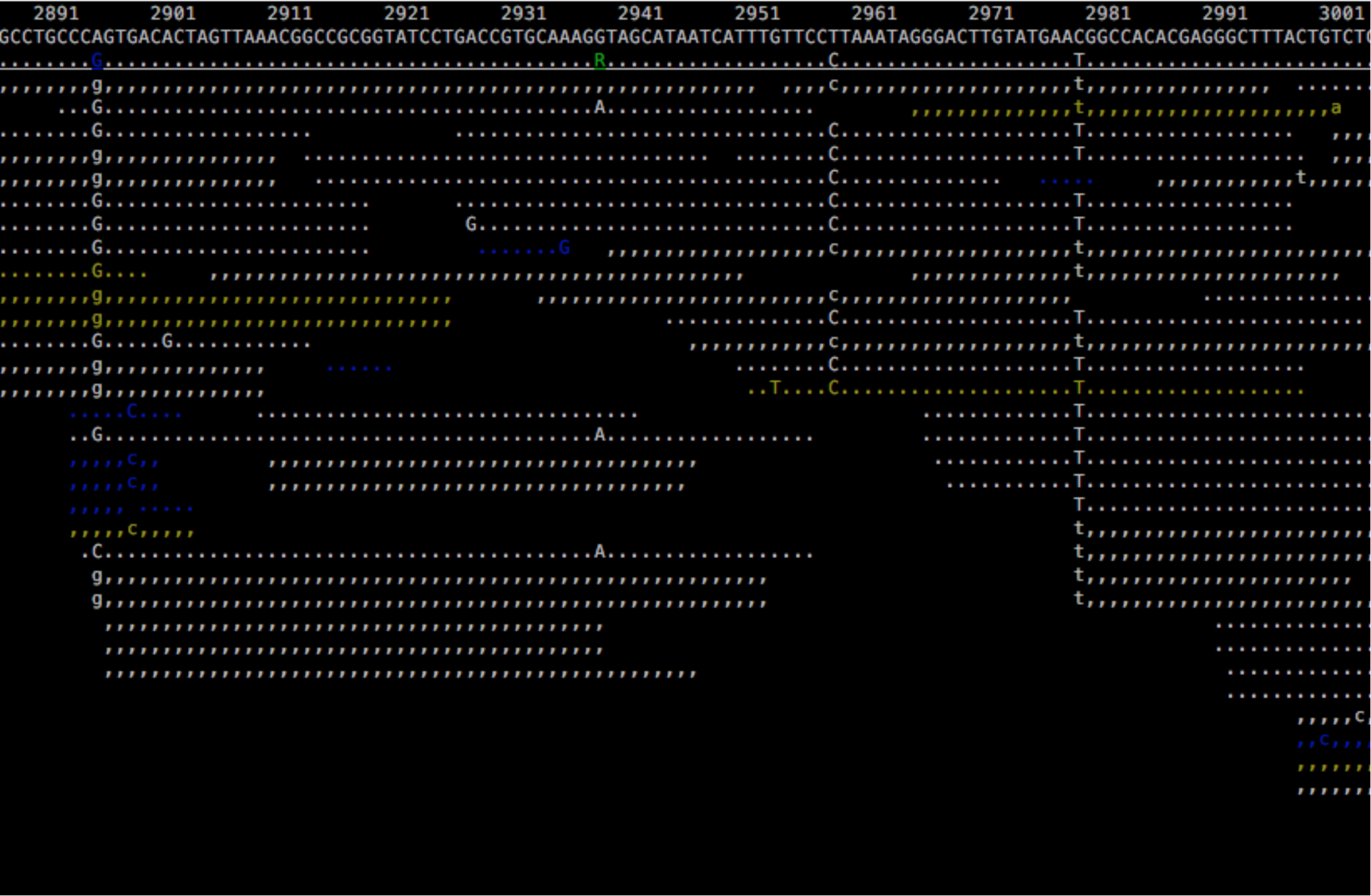
Example visualization of sequence reads and SNPs. Alignment visualization using the SAMtools package. The reference *P. concolor* mtDNA genome sequence is shown in the top sequence line, and the consensus sequence for our sample Wmt813 is shown on the second line, with individual aligned reads below. Wmt813 positions that are identical to the reference sequence are shown as “.” and “,” and colors for the aligned reads reflect sequence quality, with white being highest and blue lowest. SNPs (sequence differences between the reference sequence and Wmt813) are located at positions 2896, 2959, and 2980.

To verify that numts (nuclear mitochondrial DNA sequences) were not incorporated into our consensus sequences, we used the MITSO WebServer^31^ to annotate our sequences and translated the 13 protein-coding genes using MEGA7^32^. We found no evidence of premature stop codons or frameshift mutations. Searches were conducted using MegaBLAST^33^ using our novel sequences as queries against the mitochondrial and nuclear sequences in the Felidae family. This analysis revealed high sequence homology between the mitochondrial sequences and low sequence homology when compared to the nuclear sequences, indicating that our consensus sequences are not likely to be numts. This result is expected, given the higher per-cell copy number of mitochondrial vs. nuclear genome sequences.

### Multi-sequence alignment and phylogenetic analysis

We compared our five complete mtDNA genome sequences to previously published mitochondrial DNA sequences for modern populations of mountain lions in the Western U.S. and Florida^3,16,34^. Outside of the complete mtDNA genome sequences newly determined in this study, the majority of available (published) mtDNA sequences consisted of only part of the *P. concolor* mtDNA genome. Specifically, the majority of our analyses were necessarily based on analyses of multi-sequence alignments of only the 16S rRNA, ATPase8, and NADH dehydrogenase subunit 5 (ND5) gene regions. Sequences were aligned using MAFFT7^35^ default settings with the E-INS-i iterative refinement method, and verified visually. Multi-sequence alignment files are available on ScholarSphere: https://scholarsphere.psu.edu/collections/3484zh052. Genetic distance and phylogenetic tree analyses were conducted with MEGA7^36^. Genetic distance was calculated using the absolute number of base pair differences (p-distance) and the Kimura 2-parameter model^37^. The best substitution method was determined using jModelTest^38,39^. Phylogenetic trees were constructed using the maximum-likelihood method with 500 bootstrap replicates and the Hasegawa-Kishino-Yano model^36,40^. PopArt 1.7 (available at http://popart.otago.ac.nz) was used to generate a haplotype network with the Median-Joining Network method. All positions containing gaps and missing data were eliminated for all analyses.

## Results

We reconstructed complete mtDNA genome sequence for five of the six historical mountain lion samples that we obtained from museums and colleges in Pennsylvania. The only other currently available mountain lion complete mtDNA genomes are the *P. concolor* reference sequence^16^, which was obtained from a Texas mountain lion, and the *P. concolor coryi* sequence^34^, obtained from a Florida mountain lion (**Supplemental Table 2)**. Across 14,970 aligned bp, 4 positions were variable among the five Northeastern mountain lion individuals, 32 positions differed between all five Northeastern mountain lions and the Texas individual, 4 positions differed between Florida and the Northeast, and 36 positions were distinct between Florida and Texas. The three populations had an overall mean genetic distance (p-distance and Kimura 2-parameter model) of 0.0008 (standard error 0.0001). Pairwise distances among the Northeastern mountain lion complete mtDNA genomes, the Florida mountain lion mtDNA genome, and the reference sequence are reported in **Table 4**. Phylogenetic relationships were determined using the maximum-likelihood method, with all Northeastern mtDNA genomes and the Florida mtDNA genome clustered to the exclusion of the Texas reference mtDNA genome (**Figure 4**).

**Table 4:** **Estimates of Nucleotide Sequence Distance Among U.S. Mountain Lions**

|  | *Puma concolor*<br>reference | *P.*<br>*concolor*<br>*coryi*<br>(Florida) | Alb813 | BE813 | Har813 | RSL813 | Wmt813 |
| --- | --- | --- | --- | --- | --- | --- | --- |
| *Puma concolor*<br>reference (Texas) |  | 0.0004 | 0.0004 | 0.0003 | 0.0004 | 0.0004 | 0.0004 |
| *P. concolor coryi*<br>(Florida) | 0.0027 |  | 0.0001 | 0.0001 | 0.0001 | 0.0001 | 0.0001 |
| Alb813 | 0.0024 | 0.0003 |  | 0.0001 | 0.0001 | 0.0000 | 0.0000 |
| BE813 | 0.0023 | 0.0005 | 0.0003 |  | 0.0001 | 0.0001 | 0.0001 |
| Har813 | 0.0023 | 0.0003 | 0.0001 | 0.0003 |  | 0.0001 | 0.0001 |
| RSL813 | 0.0024 | 0.0003 | 0.0000 | 0.0003 | 0.0001 |  | 0.0000 |
| Wmt813 | 0.0024 | 0.0003 | 0.0000 | 0.0003 | 0.0001 | 0.0000 |  |

The number of base substitutions per site from between sequences are shown. Standard error estimates are shown above the diagonal and were obtained by a bootstrap procedure (500 replicates). Analyses were conducted using the Kimura 2-parameter model<sup>37</sup>. The analysis involved 7 nucleotide sequences. Codon positions included were 1st+2nd+3rd+Noncoding. All positions containing gaps and missing data were eliminated. There were a total of 14970 positions in the final dataset.

**Figure 4:**
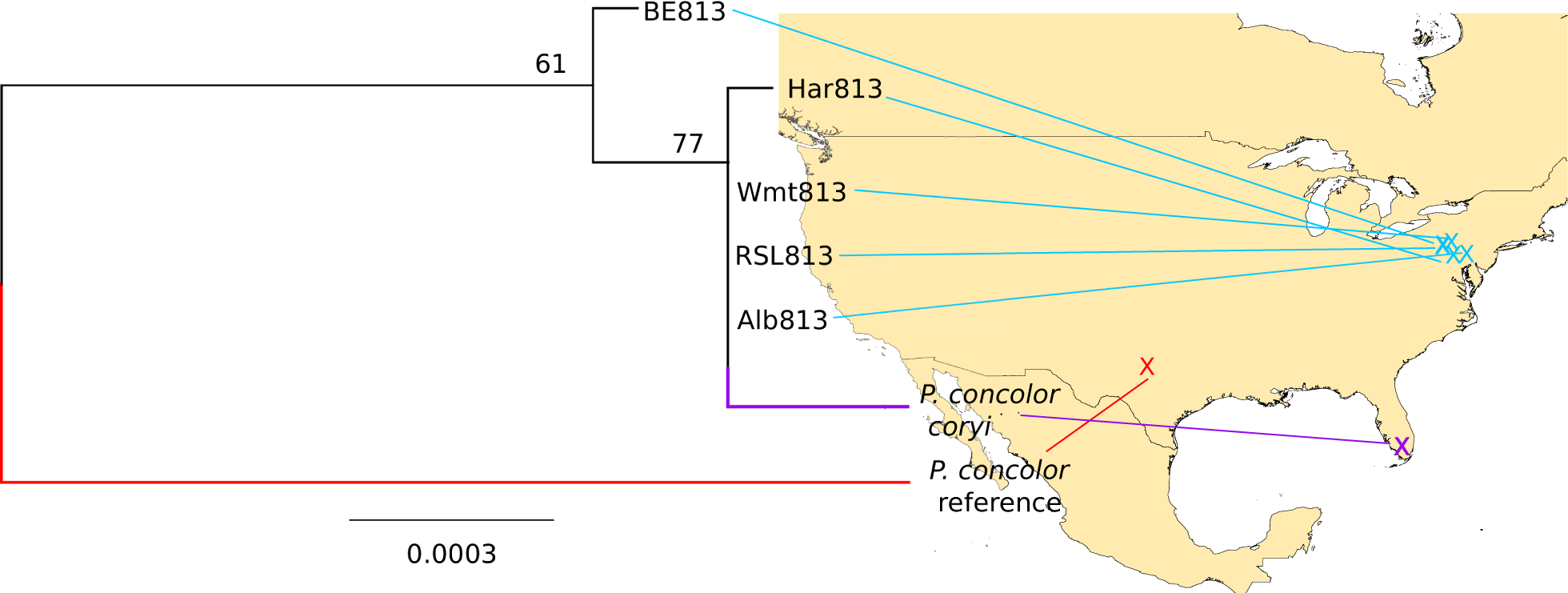
Phylogenetic analysis of six complete *P. concolor* mitochondrial genomes. The evolutionary history was inferred by using the Maximum Likelihood method based on the Hasegawa-Kishino-Yano model^50^. The tree is drawn to scale, with branch lengths reflecting the number of substitutions per site. Nodes of the ML tree are a consensus of 500 replicates. The tree with the highest log likelihood is shown (-log likelihood = 20498.91). Bootstrap values (500 replicates) are shown next to the branches. All positions containing gaps and missing data were eliminated. A total of 14970 positions were analyzed. Map source: ESRI. The map was generated using ArcGIS Pro 1.2 by ESRI (available at: http://www.esri.com/).

We also compared our sequences and the complete mitochondrial genomes from Texas and Florida to a dataset from Culver et al.^3^ that contains a large number of mountain lion individuals from North and South America (Supplemental Table 3; n=286 individuals), but only a small subset of the mitochondrial genome (the 16s rRNA, ATPase8, and ND5 genes). Among all North American mountain lions in our combined dataset (n=192), three total single nucleotide polymorphisms (SNP) were observed across the 796 bp available for analysis, representing three distinct haplotypes (**Table 5; Figure 5**). Haplotype two (H2) was common to all three populations included in this analysis (Florida panthers, Western mountain lions, and Northeastern cougars), while H1 and H3 were specific to samples from Texas and Washington State respectively.

**Table 5:** **Geographic Location and Distribution of Haplotypes in U.S. Mountain Lions**

|  | 16s rRNA<sup>a</sup> |  | ND5 |  |  |
| --- | --- | --- | --- | --- | --- |
| Haplotype | 2959<sup>a</sup> | 2980<sup>a</sup> | 12955<sup>a</sup> | Number of<br>individuals | Geographic<br>Location |
| H1 | T | C | T | 1 | Texas |
| H2 | C | T | T | 187 | Western U.S.,<br>Northeastern<br>U.S., Florida |
| H3 | C | T | C | 4 | Washington State |

<sup>a</sup> Positions based on the *Puma concolor* reference mtDNA sequence<sup>16</sup>

**Figure 5:**
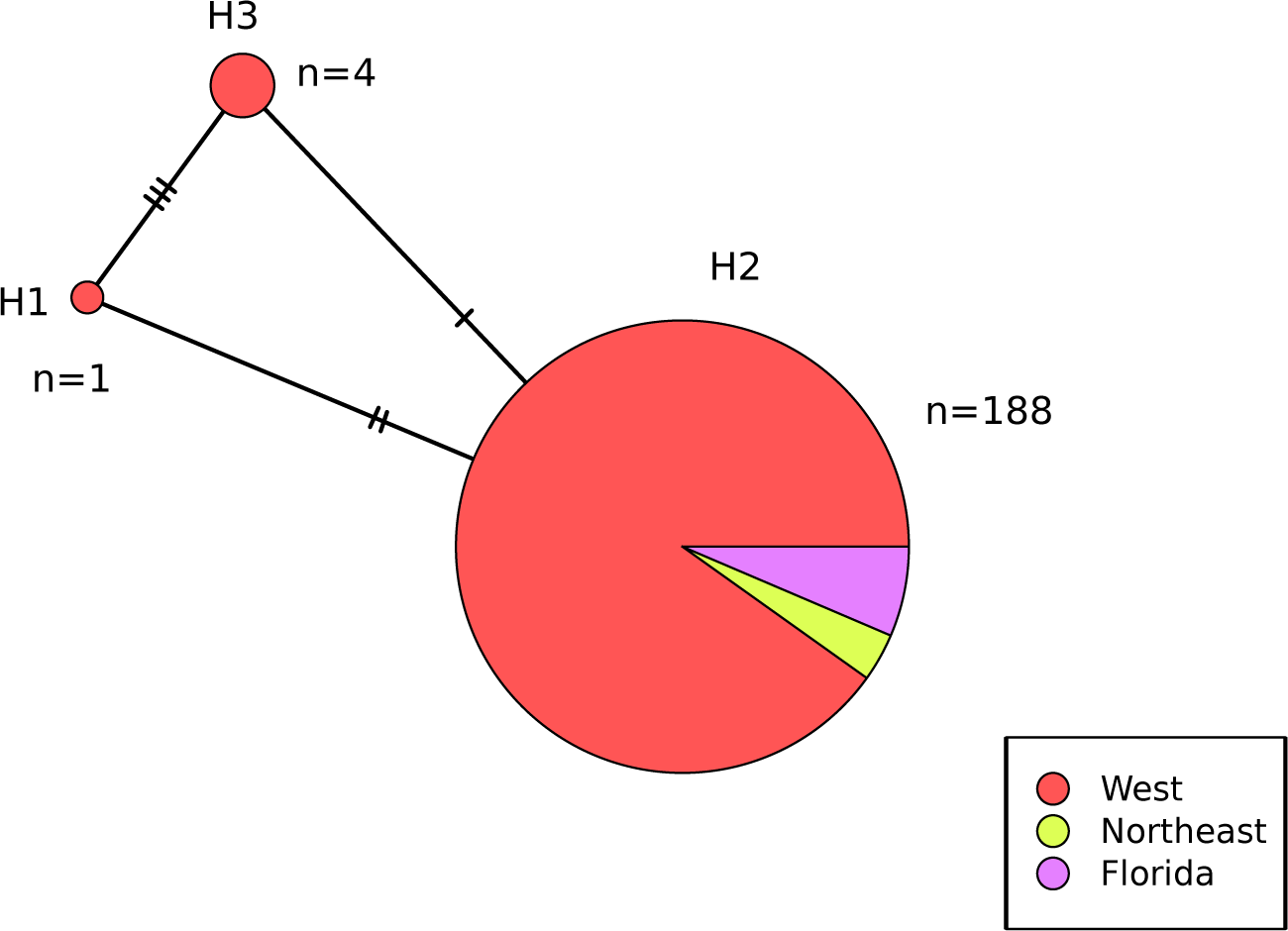
Median-joining Network Based on 3 Gene Regions for U.S. Mountain Lions. Haplotype network was generated using PopArt 1.7 (available at http://popart.otago.ac.nz) and the Median-Joining Network method. A total of 193 individuals and 796 aligned nucleotides were analyzed. All positions containing gaps and missing data were eliminated. The area of each circle is proportional to the number of samples in each haplotype. Each circle represents a single haplotype and each color represents a population. Hatch marks represent the number of single nucleotide differences between each haplotype.

We next incorporated into our analysis the 100 Central and South American mountain lion individuals with 16s rRNA, ATPase8, and ND5 mtDNA gene sequence data provided by Culver et al.^3^. All South American mountain lion mtDNA sequences are more closely related to each other than they are to North American mountain lions, and vice versa (**Figure 6**). In Central America, haplotypes identical to or similar to those from both Central America and North America are represented.

**Figure 6:**
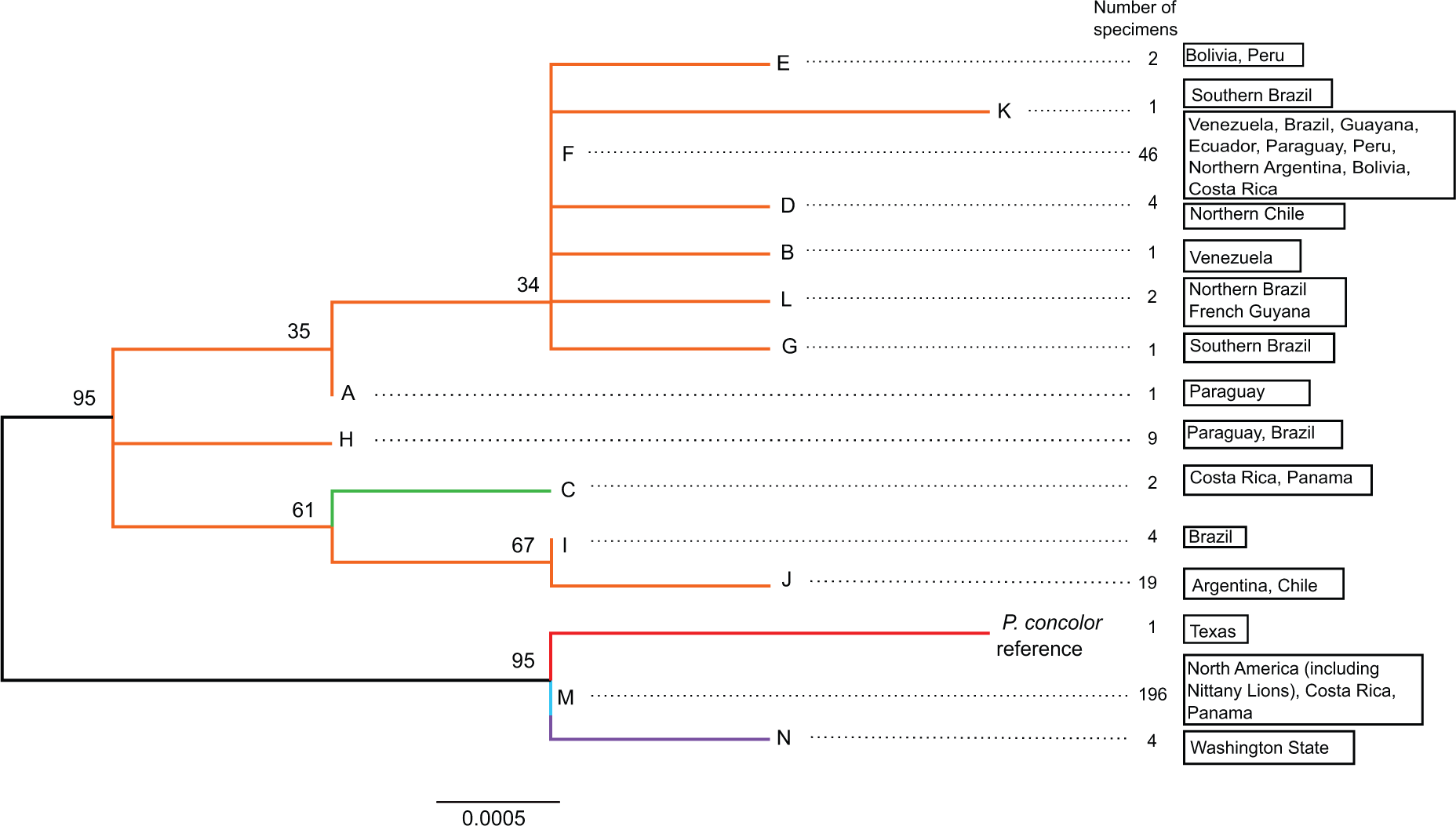
Phylogenetic analysis of three mtDNA gene regions for North, Central, and South American mountain lions. The evolutionary history was inferred by using the Maximum Likelihood method based on the Hasegawa-Kishino-Yano model^50^. Nodes of the ML tree are a consensus of 500 replicates. The tree with the highest log likelihood (-1215.29) is shown. The tree is drawn to scale, with branch lengths reflecting the number of substitutions per site. Bootstrap values (500 replicates) are shown next to the branches. All positions within the three mtDNA gene regions analyzed (16s rRNA, ATPase8, and ND5) with gaps and missing data were eliminated; there were a total of 796 positions in the final dataset. The letters shown at the end of each branch reflect the haplotype groups used by Culver et al.^3^ The five Nittany Lion samples from this study are haplotype group M.

## Discussion

This study supports previous findings that mountain lion populations from the western United States, Florida, and the Northeastern United States (extinct) are not differentiated based on their mitochondrial genomes, and that mtDNA genome diversity among North American mountain lions is and was historically relatively low overall^4,41,42^. In the past century, Western mountain lions have occasionally travelled several hundred miles to the Eastern U.S.^13,14^ While these migrating individuals are typically male (and mtDNA is inherited through the female lineage), these events underscore the large dispersal distances (males: 31-100 miles; females: 18 miles)^7^ and home range sizes (males: 25-500 square miles; females: 8-400 square miles)^6^ of mountain lions, which may have helped to limit levels of genetic differentiation across the United States^13,14,43–45^.

However, this picture may change with the future availability of complete mtDNA genomes for more Western U.S. mountain lions than the single reference individual from Texas, and especially with the availability of sequence data from the nuclear genome. Most of the nuclear genome is inherited from both parents, and at ~3 Gb (3 billion bp) the felid nuclear genome is considerably larger than the mtDNA genome. While no *Puma concolor* nuclear genome sequences have yet been published, as a follow-up to this project we next plan to generate nuclear genome sequence data for at least two of the Nittany Lion individuals from this study (particularly those with the highest endogenous DNA content; BE813 and RSL813). Analyses of these data will be incorporated into undergraduate bioinformatics classroom opportunities for students at Penn State. When ultimately combined with similar data for Western and Florida mountain lions, this work can benefit future conservation efforts and potentially help provide insight into how mountain lions have adapted to their various habitats across the continent.

Although only Florida panthers are currently listed on the endangered species list by the U.S. Fish and Wildlife Service, all mountain lions are important members of the ecosystems they inhabit^43,46,47^. The loss of this apex predator from the forests of the Northeastern U.S.A. likely had downstream consequences, including deer overpopulation^8,47,48^. Mountain lions may eventually make a return to the Northeast via the slow expansion of Western and Midwestern populations^13,49^. Our work will ultimately characterize the genetic diversity that was permanently lost (if any) from this region with the original extinction, and we hope that our associated outreach activities in the Pennsylvania region will help to raise awareness about the importance of broader ecosystem diversity in advance of the likely return of mountain lions to the Northeast.

## Acknowledgments

This project was crowdfunded, with more than 140 generous individual donors and additional support from the Penn State College of the Liberal Arts Undergraduate Research Fund, the Mountain Lion Foundation, Felidae Conservation Fund, and the Penn State Huck Institutes of the Life Sciences. Sequencing materials and DNA capture reagents used for this study were kindly donated by Illumina, Inc. and MYcroarray, respectively. The Nittany Lion specimens were provided by the Penn State All-Sports Museum, the Albright College Biology Dept., the Thomas T. Taber Museum of the Lycoming County Historical Society, Bill and Carol Thomas, and the Pennsylvania State Museum. We thank Craig Praul and the Penn State Huck Institutes of the Life Sciences Genomics Core Facility for performing the Illumina sequencing and for their support. The computational resource instrumentation used in this study was funded by the National Science Foundation (OCI–0821527). We thank Rose Carter from the Penn State University Schreyer Honors College for her enthusiasm and support of the project, and Logan Kistler, Alexis Sullivan, Kate Thompson, David Villalta, Richard Bankoff, Kyle Hilliard, Adrijana Vukelic, Baily Hohman, Joe Orkin, and Abby Koenig of the Perry laboratory for their engagement with the project, technical support, and suggestions on an earlier version of the manuscript. The findings and conclusions in this article are those of the author(s) and do not necessarily represent the views of the U.S. Fish and Wildlife Service.

## Author contributions statement

G.H.P. and M.N.E. conceived the study. M.N.E. and S.J. collected samples. M.N.E., R.J.G., and S.J. conducted experiments and analyzed the data. S.D. helped analyze the data and prepared figures 1 and 4. M.N.E. and G.H.P. wrote the paper, with review and comments by all authors.

